# Swine influenza A virus infection dynamics and evolution in intensive pig production systems

**DOI:** 10.1101/2023.10.31.564883

**Authors:** Paula Lagan, Michael Hamil, Susan Cull, Anthony Hanrahan, Rosanna M. Wregor, Ken Lemon

**Affiliations:** Veterinary Sciences Division, Agri-Food and Biosciences Institute, Belfast, Northern Ireland; JMW Farms Ltd., Armagh, Northern Ireland; Craigavon Area Hospital, Craigavon, Northern Ireland; School of Biological Sciences, Queen’s University Belfast, Belfast, Northern Ireland

## Abstract

Swine influenza A virus is one of the main viral pathogens responsible for respiratory disease in farmed pigs. Whilst outbreaks are often epidemic in nature, increasing reports suggest that continuous, endemic infection of herds is now common. The move towards larger herd sizes and increased intensification in the commercial pig industry may promote endemic infection, however, the impact that intensification has on swine influenza A virus infection dynamics and evolution is unclear. We carried out a longitudinal surveillance study over 18 months on two endemically infected, intensive pig farms. Frequent sampling of all production stages using individual and group sampling methods was performed, followed by virological and immunological testing and whole genome sequencing. We identified weaned pigs between 4-12 weeks old as the main reservoir of swine influenza A virus on the farms, with continuous, year-round infection. Despite the continuous nature of viral circulation, infection levels were not uniform, with increasing immunity at the herd level associated with reduced viral prevalence followed by subsequent rebound infection. A single virus subtype persisted on each farm for the entire duration of the study. Viral evolution was characterised by long periods of stasis punctuated by periods of rapid change coinciding with increasing immunity within the herd. An accumulation of mutations in the surface glycoproteins consistent with antigenic drift was observed, in addition to amino acid substitutions in the internal gene products as well as reassortment exchange of internal gene segments from newly introduced strains. These data demonstrate that long-term, continuous infection of herds with a single subtype is possible and document the evolutionary mechanisms utilised to achieve this.

**Author Summary:** Infection with influenza A virus is widespread in pigs and contributes towards poor health and loss of productivity. Effective infection control measures are necessary to limit the impact of influenza on farms. However, modern farming practices are increasingly characterised by larger herd sizes and higher stocking densities. It is unclear how the move towards increased intensification impacts infectious diseases such as influenza. Accumulating evidence suggests that long-term infection of herds may be common. How influenza A virus can persist long-term is poorly understood. The aim of the current study was to monitor the infection status of pigs at each stage of the production process, measure the extent of the developing immune response and to characterise the evolutionary changes in the circulating influenza A viruses over time. Our findings provide insights into the dynamics of viral infection and evolution and may contribute to improved infection control strategies.

## Introduction

Swine influenza A virus (swIAV) is a major pathogen of pigs. In naïve animals, infection usually presents as an acute respiratory disease characterised by high morbidity and low mortality [1]. Primary swIAV infection can lead to pneumonia caused by opportunistic secondary pathogens and along with Porcine Reproductive and Respiratory Syndrome virus (PRRSv), is one of the main viral factors in the development of Porcine Respiratory Disease Complex (PRDC) [2]. PRDC is a leading health concern for producers globally and has significant economic impact. Efforts to control the spread of swIAV are mostly limited to management practices that reduce mixing of animals from different sources, such as all in/all out, and vaccination of pregnant sows to protect piglets via maternal derived antibodies (MDA).

Immunity to swIAV is primarily mediated by the development of neutralising antibodies to the surface glycoproteins, hemagglutinin (HA) and neuraminidase (NA). For H1 subtypes, the major antigenic sites have been mapped to five regions on the globular head of HA, designated Sa, Sb, Ca1, Ca2 and Cb [3,4]. Although less studied, key antigenic regions on NA have also been identified [5–7]. Substitutions at antigenic sites in HA and NA leads to increased antigenic distances over time (antigenic drift), facilitating reinfection of previously immune hosts. The segmented nature of the swIAV genome can also result in reassortment of genetic segments during co-infection with two or more strains (antigenic shift). Indeed, pigs have long been hypothesised to be important in the wider ecology of influenza A virus (IAV), acting as a mixing-vessel for reassortment of avian, swine and human-adapted strains, due to the presence of both avian-like and human-like IAV receptors in the porcine respiratory tract [8,9].

Influenza is endemic in European swine, with a recent large-scale surveillance study estimating prevalence at 56.6% at the herd level [10]. The evolutionary history of swIAV is complex. Like the situation in humans, H1 and H3 subtypes with different combinations of N1 and N2 circulate globally, however, the origin, antigenicity and internal gene constellations are markedly different between North American, European and Asian lineages. In Europe, classical swine H1N1, a descendent of the virus that emerged in humans in 1918, circulated exclusively until the 1970s [11]. In 1979, classical H1N1 was replaced by an avian-like H1N1 (avH1N1), which continues to circulate and is the most frequently detected lineage in European swine herds [10,12]. A seasonal human H3N2 strain entered the European swine population in 1984 and reassorted with avH1N1 to produce swine H3N2 (swH3N2) [13]. A decade later, human-like H1N2 (huH1N2) emerged as a result of reassortment between human seasonal H1N1, swH3N2 and avH1N1 [14]. With the emergence of pandemic H1N1 in 2009 (pH1N1), reverse zoonosis into pigs occurred and pH1N1 strains are now widespread [15]. In addition, pH1N1 have contributed gene segments to other contemporary lineages via reassortment [16]. The diversification and evolution of swIAV lineages therefore represents a significant challenge to efforts that aim to control influenza via vaccination.

Under the European Union Integrated Pollution Prevention and Controls (IPPC) Directive (https://eur-lex.europa.eu/eli/dir/2008/1/oj), pig farms with 750 or more sows or 2,000 or more production pigs over 30kg are considered intensive. In Northern Ireland, as in the rest of the United Kingdom, the pork industry has moved towards increased intensification over the last two decades. Agricultural census data for 2022 shows that 87.5% of the 738,540 pigs registered in Northern Ireland were present on only 73 intensive farms (https://www.daera-ni.gov.uk/sites/default/files/publications/daera/Agricultural%20Census%202022%20Publication_1.pdf). Intensification presents a challenge for the control of infectious diseases such as swIAV. Under traditional, non-intensive pig rearing systems, swIAV outbreaks followed a typical epizootic pattern, with onset of clinical signs throughout the herd, followed by rapid recovery and the development of strain-specific herd immunity [1]. However, the move to intensification may promote long-term infection of herds, due to high turnover rates and the continuous introduction of swIAV antibody naïve animals maintaining a large population of susceptible pigs in which the virus can circulate continuously. Modelling based estimates of the minimum population size necessary for continuous infection are relatively modest, at around 3,000 pigs [17]. Cohort studies (following batches of pigs from birth to slaughter or through specific production stages) and longitudinal studies, (where specific populations of animals are sampled repeatedly over time) have provided valuable insights into swIAV infection dynamics in a variety of production systems [18–27]. However, the impact of intensification on swIAV infection dynamics and evolution is unclear. We therefore carried out longitudinal surveillance on two intensive swine farms over an 18-month period in order to understand (i) which production stage(s) are most susceptible to infection, (ii) whether infection is epizootic or enzootic in nature and (iii) if maintaining a large population of swIAV antibody naïve animals lessens selection pressures that normally drive antigenic drift and shift.

## Materials & Methods

### Farm selection, organisation and vaccination status

Three multi-stage, intensive commercial pig farms located in Northern Ireland were visited in January 2021 and tested for the presence of swIAV by real-time reverse-transcription polymerase chain reaction (RRT-PCR). All production stages including suckling pigs (sucklers), weaned pigs (weaners), finishing pigs (finishers) and gilts were sampled using nasal wipes, udder wipes and oral fluid sampling. A reduced number of samples were collected during this initial visit compared to subsequent visits (46 versus 55 samples per farm). Two of the three farms tested positive for swIAV by RRT-PCR in at least one production stage and were selected for inclusion in the longitudinal study, with a total of 12 visits over 18 months.

The organisation and herd details for each farm are shown in Fig 1. Lactating sows were kept in farrowing crates. Piglets remained with the sows until four weeks of age, after which they were transferred to the nursery facilities on the weaning units. The weaning units had first and second stages (i.e.,7kg-15kg and 15kg-30kg). Pigs remained on the weaning unit until they reached 30kg at around 12 weeks old, after which they were transported to the finishing facilities where they remained until reaching slaughter weight at approximately 24 weeks. Breeding, weaning and finishing sites were all geographically separate and had temperature and ventilation controls. Gilts, weaners and finishers on the three units were housed in pens across multiple rooms. Replacement gilts were obtained from a separate nucleus herd and were brought onto each individual breeding site from 4 to 8 weeks old every month. For both farms, management practices included the use of all in/all out at the room level and vaccination of gilts and pregnant sows for influenza A virus. Gilts received inactivated tri-valent swIAV vaccine (Respiporc FLU3, IDT Biologika) at 6 and 3 weeks before service. Sows were vaccinated 3 weeks prior to farrowing.

**Fig 1.**
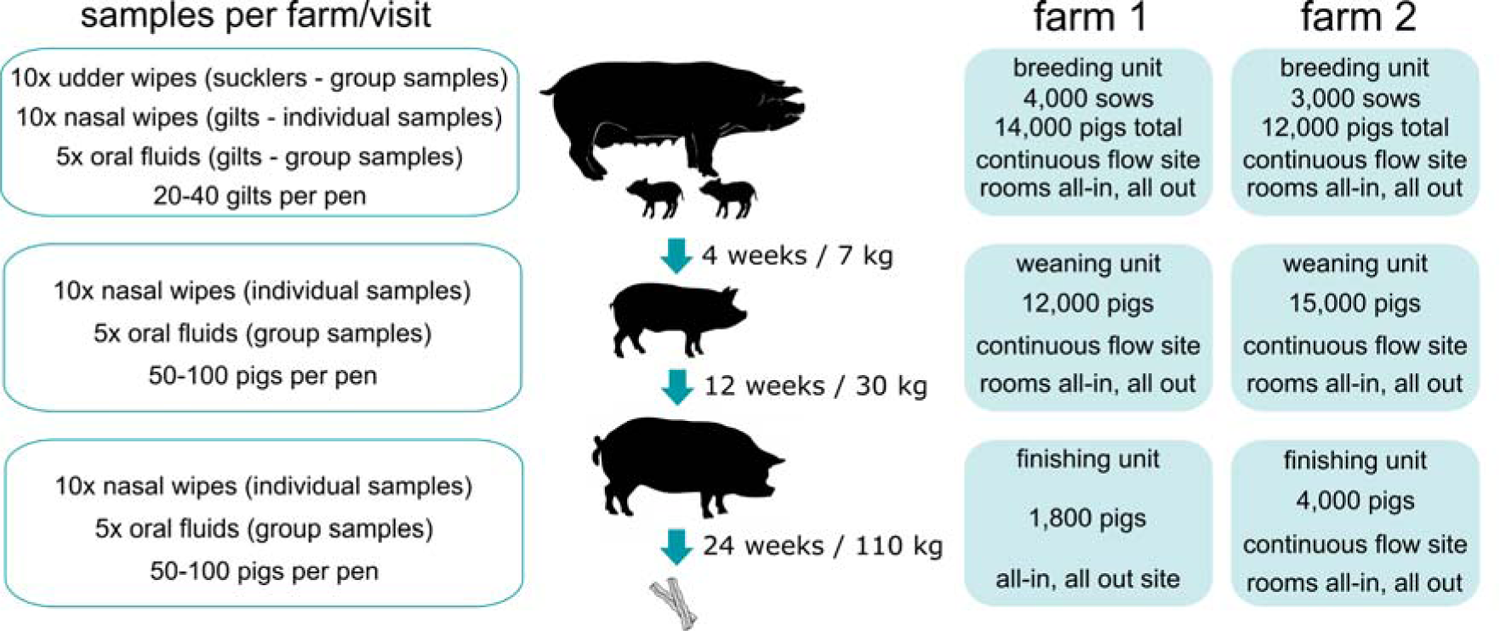
Overview of sampling strategy, farm organisation and management practices. (Left) Sample types and number of samples collected per production stage per visit. (Middle) Production stages showing progression milestones from farrow to slaughter. (Right) Farm organisation including population at each production stage and management practices.

### Ethical statement

In this study, only non-invasive samples (nasal wipes, udder wipes and oral fluids) were collected and therefore did not require a specific licence according to UK Law (Animals Scientific Procedures Act 1986). A trained veterinarian was involved in the sampling, and consent of the farmer was obtained before sample collection.

### Sample collection & processing

An overview of the sampling strategy used in this study is shown in Fig 1. Ten nasal wipes each were collected from gilts, weaned pigs and finishing pigs per farm per visit. Nasal wipes were collected by wiping a sterile 5cm x 5cm cotton gauze (MediSupplies Ltd. #PMC5771) across the pig’s snout which was placed into a sterile Nalgene Wide-Mouth HDPE 30ml Bottle (ThermoFisher Scientific #2104-0001) containing 5ml Brain-Heart Infusion Broth (Merck #53286) supplemented with antibiotics/antimycotics (1000IU penicillin G; 10µg/ml amphotericin B; 1mg/ml gentamicin). Ten udder wipes were collected from sows/suckling pigs per farm per visit using the same reagents as used for nasal wipes. A single cotton gauze was used to collect the nasal/oral secretions deposited on the udders by the litter during or immediately after suckling and placed in bottles containing transport media as above. Oral fluids were collected using Oral Fluid Collection Kits (IDEXX #98-0002094-00). Undyed cotton, 3-strand twisted ropes were placed hanging from pens and pigs allowed to chew for approximately 30mins. Ropes were squeezed into plastic bags and the recovered fluids transferred to 15ml centrifuge tubes. A single oral fluid sample was collected per pen, with 5 pens each sampled for gilts, weaned pigs and finishing pigs per farm per visit.

For all sample types, gloves were worn and changed between each sample collection. Pens, litters or individual pigs were selected whenever possible based on the presence of clinical signs suggestive of swIAV infection, such as coughing or sneezing, and randomly when clinical signs were not observed. Following collection, samples were stored at 2-8°C for up to 4 days before transport to the laboratory. Upon receipt, nasal/udder wipes (both gauze and unabsorbed transport media) were transferred to 10ml centrifuge tubes and centrifuged at 2683g for 10min at 4°C. Following centrifugation, media present above the pelleted gauze was collected, aliquoted and stored at −80°C. Oral fluid samples were centrifuged at 2683g for 10min at 4°C to remove debris and supernatants aliquoted and stored at −80°C.

### RNA extraction & RRT-PCR

Nucleic acids were extracted from 0.2ml of clinical material or allantoic fluid using the IndiMag Pathogen Kit (Indical Bioscience #SP947457) on a KingFisher Flex purification system with 96 deep-well head (ThermoFisher Scientific #5400630) according to the manufacturers’ instructions. Extracted material was tested for the presence of influenza virus RNA by RRT-PCR using primers and probe which detect a conserved region in the viral matrix gene, as described previously [28]. RRT-PCR reactions were ran using AgPath-ID One Step RT-PCR Reagents (ThermoFisher Scientific #4387391) on a LightCycler 480 (Roche). Two replicates per sample were tested, with samples returning threshold cycle (ct) values <38 in duplicate considered positive.

### Influenza anti-NP blocking ELISA

The detection of swIAV anti-NP antibodies in oral fluid samples using a modified, commercially available ELISA (Influenza A Ab Test, IDEXX #99531-01) has been described previously [29,30]. Modifications to the manufacturer’s instructions include increasing the volume of sample loaded to 0.2ml and incubating plates at 21°C for 16h. Following incubation, optical densities were measured at 650nm and sample-to-negative (S/N) ratios calculated according to the manufacturer’s instructions. Two replicates were tested for each sample and the percentage coefficient variation (%CV) calculated as instructed. Samples with %CV >15 were repeated. Repeat samples returning %CV >15 were considered invalid and excluded. Valid samples with mean S/N ratios ≤0.6 were considered positive.

### Virus isolation

Ten-day old embryonated specific pathogen free chicken eggs (VALO Biomedia) were used to isolate swIAV from RRT-PCR positive clinical samples. 0.1ml of undiluted nasal/udder wipe fluid was inoculated into the allantoic cavity using a 1ml syringe fitted with a 26G needle. Oral fluid samples were supplemented with antibiotics/antimycotics (1000IU penicillin G; 10µg/ml amphotericin B; 1mg/ml gentamicin) prior to inoculation. Three eggs per sample were inoculated and allantoic fluid harvested and pooled following 4 days incubation at 37°C. Debris was pelleted at 800g for 10mins at 4°C and aliquots stored at −80°C. Successful virus isolation was confirmed either by haemagglutination of 0.5% chicken blood cells (Envigo #S.B-0008) or by RRT-PCR.

### Influenza whole genome sequencing

The selected samples for whole genome sequencing (WGS) were extracted using the QIAamp® Viral RNA Mini Kit (Qiagen #52906) with the addition of carrier RNA. A multiplex RT-PCR assay was designed to amplify the whole segmented influenza RNA genome for Next Generation sequencing (NGS). The multiplex assay was composed of primers with complementary sequence to the IAV genome universal conserved termini regions [31] and additional adapted primers with sequence specific to the 3’ end complementary of PA, PB1 and PB2 genes enabling the amplification of the larger polymerase gene segments (Supplementary Table S1). The underrepresented distal ends of the amplicons in Illumina NGS coverage are due to the nature of the transposon’s double stranded DNA cleavage mechanism, however, this was resolved by concatenating overhangs of Nextera transposase adapter sequences to each primer set as either a full length or as a truncated version (Supplementary Table S1). The amplification reaction used the SuperScript™ III One-Step RT-PCR System with Platinum™ Taq DNA Polymerase (Invitrogen #12574018). The generated amplicons were assessed for correct amplicon base pair (bp) size on a 1% agarose gel. The NGS library was prepared with the Nextera XT DNA Library prep kit following manufacturer’s instructions with changes (Illumina #FC-131-1096). Initially, ≤100 bp fragments were removed by AMPure (Beckman Coulter) bead washing. The samples were diluted to 0.2ng/µl in 10 mM Tris HCl pH 8.0 and using the Illumina protocol for tagmentation and indexing. A double-sided clean-up was completed to select an index fragment size range for sequencing with the right-hand side dilution factor at 0.8 x followed by a 1.1 x dilution factor for the lefthand side. Each sample was sized by the Gel fragment analyzer instrument using the DNF-474 High Sensitivity NGS assay (Agilent Technologies #DNF-474-0500) to calculate the average fragment size (expected fragment size of ∼600bp) before quantifying by the Qubit instrument and preparing a 4nM equimolar library. An 8pM library with 10% PhiX control was sequenced on the MiSeq instrument (Illumina) with the MiSeq reagent Kit 500v2 cycle kit at 2 x 200bp as pairend reads. The quality of NGS data was screened using Trim Galore (https://www.bioinformatics.babraham.ac.uk/projects/trim_galore/) whilst selecting a high Q ≥33 score and then reviewed by the MultiQc program [32]. Pairend reads were aligned using the Burrows-Wheeler Aligner maximum exact matches (BWA-MEM) (v0.7.17-r1188) read aligner to selected indexed references genomes A/swine/Northern Ireland/2005-22/2021 (H1N2) and A/swine/Northern Ireland/2004-30/2021 (pH1N1) in SAMtools (v1.10) [33] creating a sequence alignment map (SAM) file [34]. The SAM files were indexed to binary sequence map (BAM) files, and the read depth and breadth of coverage were determined using SAMtools before visualising, generating consensus sequences and single-nucleotide polymorphisms (SNP) calling in Geneious R10 (Biomatters). WGS data are available under GISAID accession numbers EPI_ISL_18221301 to EPI_ISL_18221395.

### HA / NA protein structure homology-modelling

HA / NA protein structure homology models of representative farm 1 and farm 2 strains were generated using Swiss-Model (https://swissmodel.expasy.org/). The HA amino acid sequence of A/Northern Ireland/2012-08/2021 (pH1N1) was modelled using the H1 of A/Michigan/45/2015 (https://doi.org/10.2210/pdb7KNA/pdb) as template, with a final Global Model Quality Estimate of 0.76. The HA of A/Northern Ireland/2006-10/2021 (huH1N2) was modelled using H1 of A/Hickox/JY2/1940 (https://doi.org/10.2210/pdb6ONA/pdb) as template, with a final Global Model Quality Estimate of 0.77. The NA of A/Northern Ireland/2006-10/2021 (huH1N2) was modelled using the N2 of A/Moscow/10/1999 (https://www.wwpdb.org/pdb?id=pdb_00007u4f) as template, with a final Global Model Quality Estimate of 0.79.

### Phylogenetic analysis of swIAV

Complete coding sequences of PB2, PB1, PA, HA, NP, NA, NS1, NEP, M1 and M2 were aligned by MUSCLE in MEGA version 11.0.13 (https://www.megasoftware.net/). The evolutionary history was inferred using the Maximum Likelihood method and Tamura-Nei model, with phylogeny tested using 1,000 bootstrap replications. Clade designation was performed using the Swine H1 Clade Classification Tool at the Influenza Research Database (http://www.fludb.org).

## Results

### Utility of nasal wipes, udder wipes and oral fluids for swAIV surveillance

Effective surveillance for swIAV requires a balance between obtaining sufficient sample numbers to detect a representative cross-section of circulating strains alongside consideration of the downstream diagnostic methods to be utilised versus cost and ease of sampling. Nasal swabs are considered the gold-standard for virological detection of swIAV. However, pigs must be suitably restrained to obtain nasal swabs, making collection of large numbers labour intensive. Alternative sampling approaches for swIAV surveillance have received attention in recent years, including nasal wipes, udder wipes, environmental swabbing, aerosol sampling and oral fluids [35–37]. Of these, nasal wipes and udder wipes have been shown to perform equivalent to nasal swabs for the detection and isolation of swIAV whilst being relatively easy to obtain. Oral fluids are an increasingly popular sample type for routine viral surveillance, allowing groups of pigs to be sampled with ease. In addition to detection of shed viruses, oral fluids have also been successfully utilised to measure secreted swIAV antibodies [29,30]. However, virus isolation rates from oral fluids are generally low and obtaining samples from suckling pigs is difficult due to lack of interest in the cotton ropes. In this study, we primarily utilised nasal wipes to obtain individual samples suitable for downstream applications including virus isolation, and oral fluids to obtain group samples to expand the breadth of surveillance. Litters of suckling pigs were sampled using udder wipes, overcoming the limitations of oral fluids for this age group.

In the current study, nasal wipes and oral fluids were found to be comparable in terms of detection of swIAV by RRT-PCR (Table 1), having positivity rates of 18.1% and 17.3%, and mean ct values of 31.4 and 30.8, respectively. However, viral isolation rates in embryonated eggs were low for oral fluids (10.9%) compared to nasal wipes (52.3%). The utility of udder wipes was less clear due to the apparent low prevalence of swIAV on the breeding units under investigation. During the 18-month study period, only 7 of 240 (2.9%) udder wipes tested positive and only 1 viral isolate was obtained.

**Table 1.**
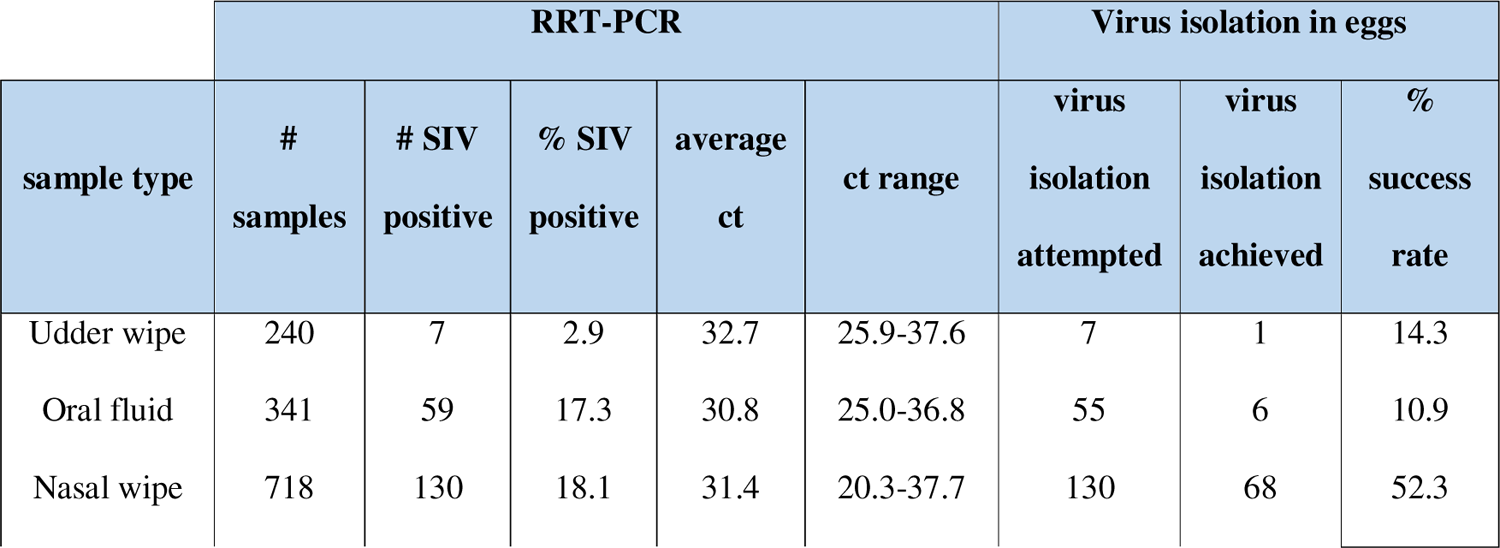
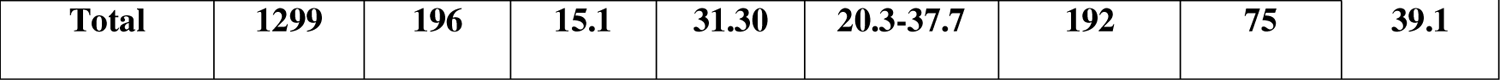
Molecular detection and viral isolation rates for swIAV from nasal wipes, udder wipes and oral fluids.

| sample type | RRT-PCR |  |  |  |  | Virus isolation in eggs |  |  |
| --- | --- | --- | --- | --- | --- | --- | --- | --- |
|  | # samples | # SIV positive | % SIV positive | average ct | ct range | virus isolation attempted | virus isolation achieved | % success rate |
| Udder wipe | 240 | 7 | 2.9 | 32.7 | 25.9-37.6 | 7 | 1 | 14.3 |
| Oral fluid | 341 | 59 | 17.3 | 30.8 | 25.0-36.8 | 55 | 6 | 10.9 |
| Nasal wipe | 718 | 130 | 18.1 | 31.4 | 20.3-37.7 | 130 | 68 | 52.3 |
| Total | 1299 | 196 | 15.1 | 31.30 | 20.3-37.7 | 192 | 75 | 39.1 |

### Prevalence of swIAV in different production stages

Fig 2 summarises the proportion of samples testing positive for swIAV by RRT-PCR from each production stage. Associated ct values of positive samples are included in Supplementary Fig S1. On both farms, the vast majority of positive samples (183 of 196) were obtained from weaned pigs between 4-12 weeks old. Virus was detectable throughout the entire 18-month study period on the weaning units apart from a single time-point (August 2021 for farm 1 and September 2021 for farm 2). Two discrete waves of infection could be discerned. For farm 1, the first wave of infection peaked during April 2021 (86.7% positivity, mean ct 29.3), followed by a decline and eventual disappearance in August 2021. The virus re-emerged in September 2021, resulting in a second wave of infection that peaked in May 2022 (100% positivity, mean ct 26.6). A similar pattern was observed for farm 2, with the initial wave peaking during May 2021 (73.3% positivity, mean ct 29.5), followed by a subsequent decline and disappearance in September 2021. The virus re-emerged in October 2021 and persisted until the end of the study, reaching peak prevalence in July 2022 (80% positivity, mean ct 31.0).

**Fig 2.**
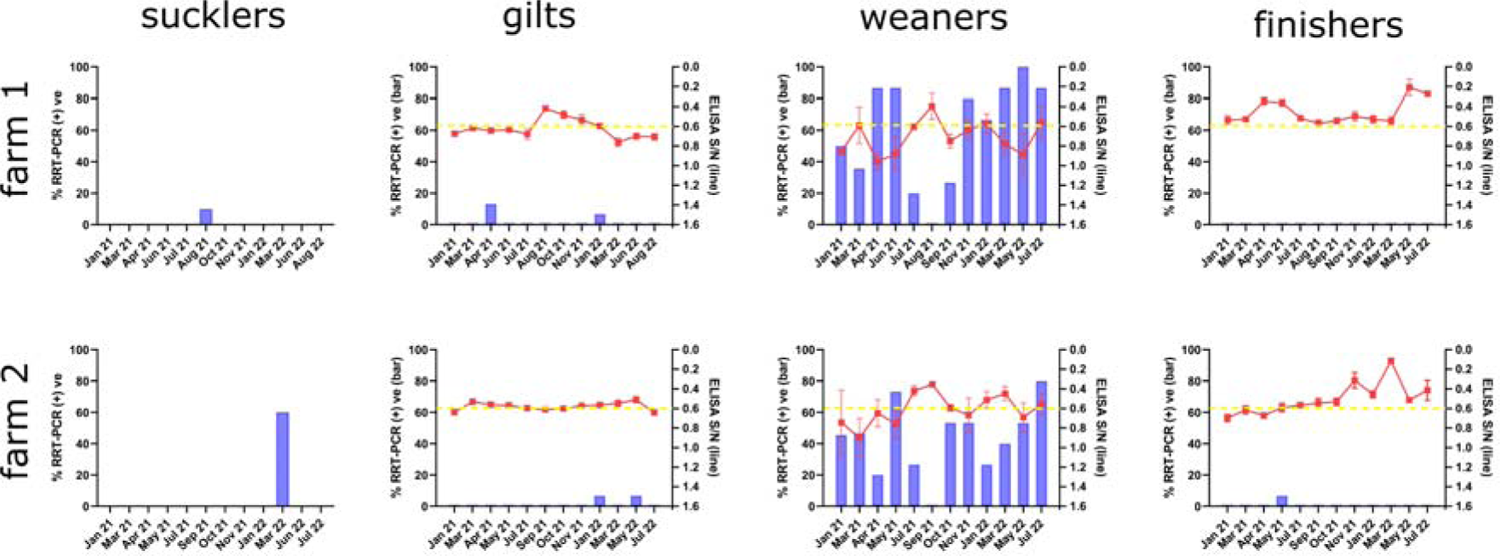
Comparison of viral prevalence with antibody status. Nasal wipes, udder wipes and oral fluids were tested for the presence of swIAV by RRT-PCR and plotted as percentage positivity (left y axis, blue bars). Each data point for gilts, weaners and finishers represents 15 samples (10 nasal wipes and 5 oral fluids), except for the initial January 2021 time-point which represents 12 samples (10 nasal wipes and 2 oral fluids). For sucklers, each data point represents 10 udder wipes (10 litters). Oral fluid samples were also tested for the presence of influenza NP antibodies by a blocking ELISA and plotted as S/N ratios on the same graph (right y axis, red line). S/N ratios ≤0.6 are considered antibody positive. The dashed yellow line indicates the 0.6 S/N ratio cut-off value.

For all other production stages, swIAV was either absent or present at a very low prevalence. Where present, detections by RRT-PCR on these units were sporadic and associated with ct values >31. An exception to this observation occurred on the farm 2 breeding unit. Following 14 months without a detection, 60% of samples tested positive for swIAV in March 2022, with ct values ranging from 25.9 - 34.9. Subsequent sampling points in June and July 2022 tested negative by RRT-PCR.

### swIAV antibody status in different production stages

Of the 341 oral fluid samples collected during this study, 11 had insufficient material remaining following molecular testing to allow testing by ELISA or failed ELISA quality control due to high %CV values of replicates and so weren’t included in subsequent analysis. ELISA data from the remaining 330 oral fluid samples analysed for the presence of swIAV specific antibodies revealed different patterns of antibody positivity for each production stage (Fig 2 and Supplementary Fig S1B). Antibody levels in gilts were relatively uniform, with ELISA S/N values trending around the assay cut-off value of 0.6 for most of the study period. A spike in antibody positivity occurred in farm 1 gilts in August 2021 followed by a return to baseline values.

Weaned pigs displayed the most variability in antibody levels, ranging from high S/N values (indicating antibody naïve pigs) to low S/N values (indicating antibody positive pigs). At the herd level, mean S/N values in oral fluid samples from weaned pigs on farm 1 inversely correlated with viral prevalence data obtained by RRT-PCR. Peak swIAV antibody levels were recorded on farm 1 during August 2021, which coincided with the disappearance of the virus between the first and second waves of infection. Conversely, the lowest antibody values were recorded in April 2021 and May 2022, coinciding with peak viral prevalence of the first and second waves respectively. For farm 2, the relationship between swIAV antibody levels and viral prevalence data was less clear. Similar to farm 1, antibody levels in farm 2 weaned pigs peaked in September 2021 and this coincided with the disappearance of the virus between the first and second waves of infection. However, a second peak in antibody levels in March 2022 occurred during a period of increasing viral prevalence.

Antibody levels in finishing pigs were characterised by long periods of uniformity around the S/N 0.6 cut-off value punctuated with spikes indicating increasing antibody levels in the herd. For farm 1 finishing pigs, these increases in antibody levels occurred during April 2021 and May 2022, whereas for farm 2 finishing pigs, antibody levels peaked in November 2021 and March 2022.

### Genetic analysis of swIAV subtypes

The WGS method employed in this study consistently produced high quality sequence data when the starting material had ct values ≤ 28. Of the 196 swIAV positive samples obtained, only 27.6% (54 of 196) were suitable for direct sequencing from the original clinical material. In order to maximise the number of viral sequences available, viral isolation in embryonated chicken eggs was attempted for all RRT-PCR positive samples, regardless of ct value. Using this approach, 75 viral isolates were obtained and complete genomic sequences determined. WGS was attempted for a subset of samples with ct values > 28 where virus isolation was unsuccessful, however these resulted in only partial genomic sequences.

Analysis of viral WGS data revealed that two swIAV subtypes were in circulation (Fig 3). For farm 1, sequence data was only obtained from weaned pigs and identified that pH1N1 clade 1A.3.3.2 was present for the entire 18-month study period (Fig 3A). During the first wave of infection, only pH1N1 was detected on the weaning unit. During the second wave, pH1N1 predominated but co-circulated with huH1N2 clade 1B.2.2 between November 2021 and May 2022. For farm 2, the majority of sequence data was obtained from weaned pigs, although a single complete genome was obtained from a suckling pig during the March 2022 outbreak in the breeding unit and a partial genome sequence was determined from a gilt in May 2022. huH1N2 clade 1B.2.2 was the dominant strain present on farm 2 and was exclusively present on the weaning unit for the entire duration of the study. The single gilt sequence was also identified as huH1N2 clade 1B.2.2. However, the strain responsible for the suckling pig outbreak on the farm 2 breeding unit was identified as pH1N1 clade 1A.3.3.2.

**Fig 3.**
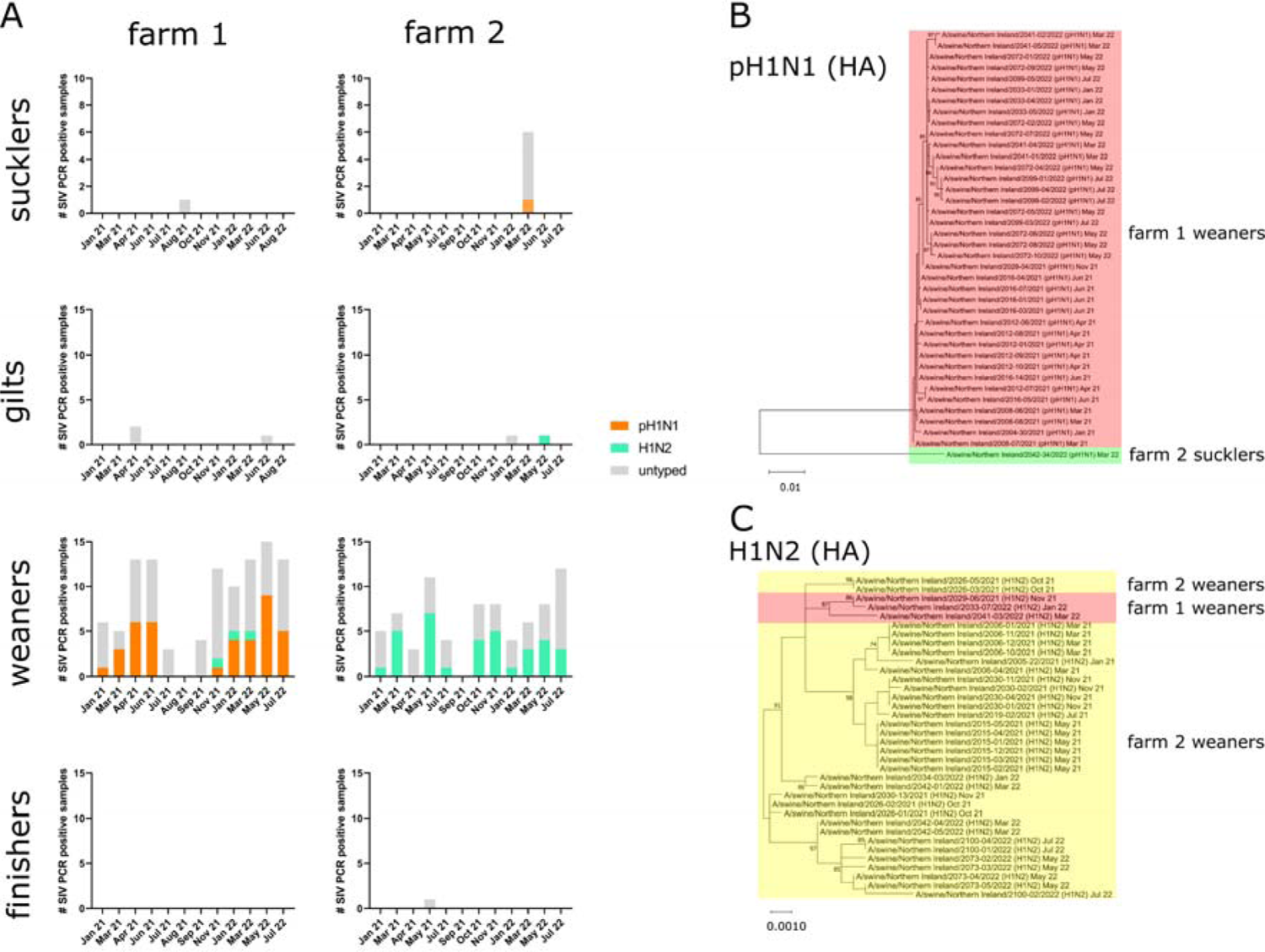
Subtyping and phylogenetic analysis of swIAV. (A) Stacked bar charts showing the proportional contribution of different swIAV subtypes to the overall prevalence for each production stage on both farms: pH1N2 (orange), huH1N2 (green) and untyped (grey). Maximum-Likelihood phylogenetic trees of all (B) pH1N1 and (C) huH1N2 sequences obtained based on complete HA coding sequences, with 1,000 bootstrap replications. Trees colour-coded according to the farm/production stage where samples were obtained: farm1 weaners (pink), farm 2 weaners (yellow) and farm 2 sucklers (green).

Phylogenetic analysis of pH1N1 HA sequences indicated that all strains detected on the farm 1 weaning unit were closely related and were distinct from the pH1N1 detected on the farm 2 breeding unit (Fig 3B). In contrast, all huH1N2 strains were closely related regardless of farm origin (Fig 3C). The 6 internal gene segments of all strains identified in this study were derived from pH1N1.

### Antigenic drift of swIAV HA and NA

As the major antigenic proteins, HA and NA accumulate amino changes in response to immune pressure, resulting in antigenic drift and immune escape. The observation of rising swIAV antibody levels on the weaning units, followed by the disappearance and subsequent re-emergence of virus, led us to speculate that increasing immunity within the herd exerted selection pressure that favoured the emergence of antigenically drifted strains. To investigate this, predicted HA and NA amino acid sequences from viral isolates obtained from the weaner units spanning the entire 18-month study period were aligned and substitutions recorded. This revealed a similar pattern of evolution over time on both farms (Fig 4). For farm 1, the pH1N1 HA and NA proteins were remarkably stable over the initial 6-month period of the first wave of infection (Fig 4A). Of the 16 strains isolated from 4 sampling time-points during this period, 12 had identical HA and NA sequences. Four strains had 1 or 2 amino acid changes in HA, all of which were only detected in a single isolate at a single time-point. With the re-emergence of the virus during the second wave of infection, several mutations in HA and NA became fixed, with strains carrying these changes displacing previously circulating strains. Mutations emerging at the start of wave 2 included S83P and V345I in HA and I10V in NA, with an additional HA S45N mutation emerging in January 2022. A variant with two additional substitutions in NA (V62I and I106V) emerged in May 2022 and co-circulated with the original wave 2 variant until the end of the study in July 2022. For farm 2, the huH1N2 HA and NA protein sequences were similarly static during the 6-month period of the initial infection wave (Fig 4D). Of the 12 strains isolated over 4 sampling time-points, most differed only at two variable positions (30 and 309) in HA. Three isolates had one or two additional substitutions in HA or NA that were only detected on one occasion. The dominant strain that emerged during the second wave of infection contained the substitutions V172I and A196T in HA and T452S in NA. A variant with an HA sequence identical to the original strain (except for a single substitution in the HA signal sequence) but with additional mutations in NA (Y40H and K415R) emerged briefly during the second wave of infection but did not persist. The dominant huH1N2 variant acquired two additional mutations in NA (V313G and K385R) in January 2022 that became fixed and displaced previous circulating strains for the reminder of the study.

**Fig 4.**
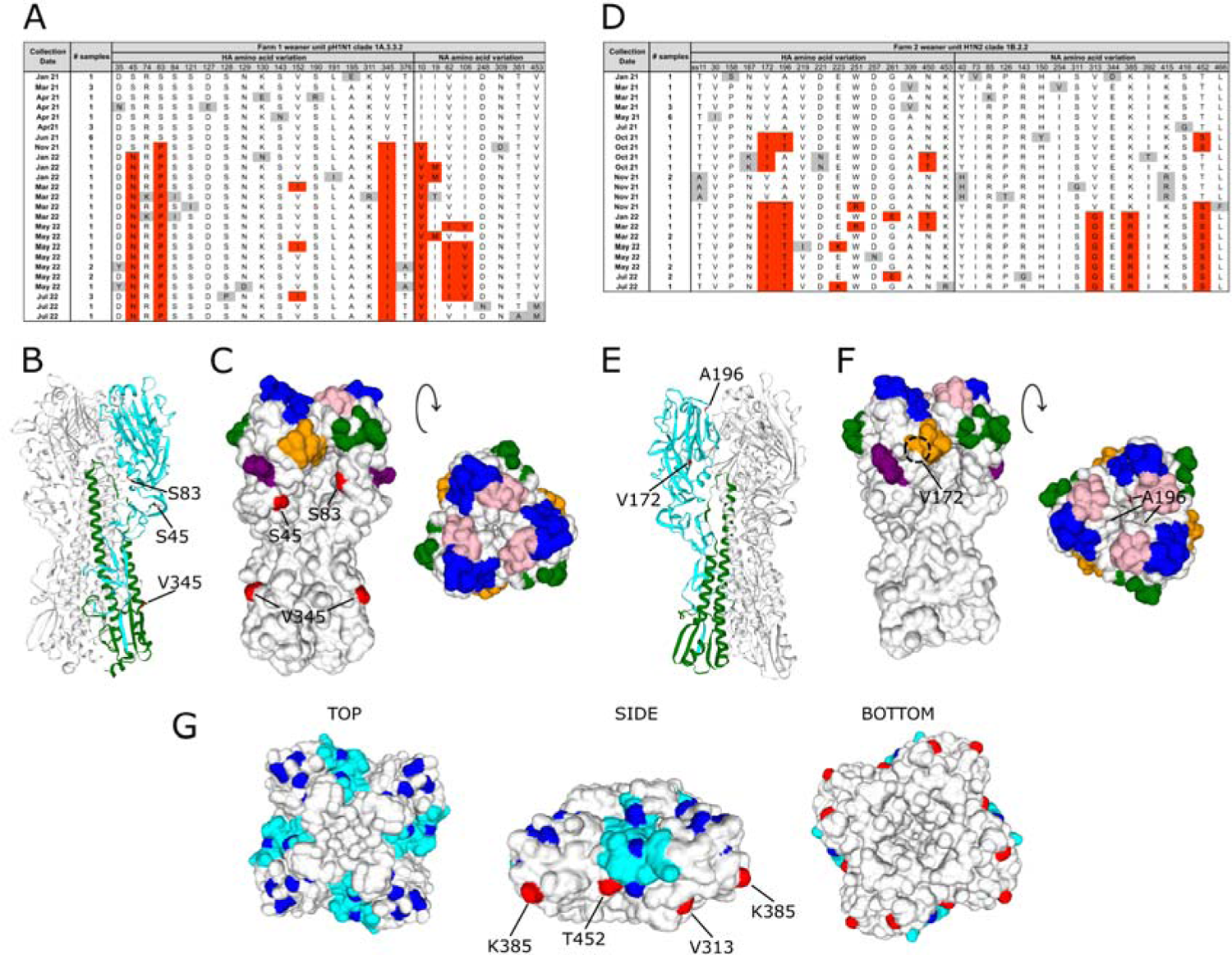
Amino acid substitutions in the swIAV surface glycoproteins over time. Amino acid substitutions in HA and NA of (A) farm 1 weaner unit pH1N1strains and (D) farm 2 weaner unit huH1N2 strains. Consensus sequences were derived from strains isolated during the first 6 months of the study. Deviations from the consensus are highlighted. Grey highlights indicate substitutions that were detected during a single time-point. Red highlights represent substitutions detected during two or more time-points. HA numbering is according to HA0, following cleavage of the 17-residue signal sequence. Substitutions occurring in the signal sequence are prefixed with ss followed by the amino acid number. HA homology models of (B) A/swine/Northern Ireland/2012-08/2021 (farm 1 weaner unit pH1N1, collection date April 2021) and (E) A/swine/Northern Ireland/2006-10/2021 (farm 2 weaner unit huH1N2, collection date March 2021). HA1 domain (cyan) and HA2 domain (green) from a single monomer is highlighted with locations of major substitutions indicated in red. Surface rendering of (C) A/swine/Northern Ireland/2012-08/2021 and (F) A/swine/Northern Ireland/2006-10/2021 HA homology models. Antigenic sites indicated: Sa (blue), Sb (pink), Ca1 (orange), Ca2 (green), Cb (purple). Locations of major substitutions highlighted in red. The location of buried residues, not visible in surface renderings, are indicated by a dashed circle. (G) NA homology model of A/swine/Northern Ireland/2006-10/2021 (huH1N2). Mem5 epitope (cyan), monoclonal antibody escape mutations (blue). Locations of major substitutions indicated in red.

Isolation of IAV in eggs can result in egg adaptation mutations, making subsequent sequence interpretation difficult [38]. In order to assess the impact of egg adaptation on the HA and NA sequences of the swIAV strains described here, WGS was performed on a subset of samples direct from original clinical material and compared with sequences from the corresponding egg isolates (Supplementary Table S2). Of the 18 paired samples sequenced (8 pH1N1 and 10 H1N2), 13 had identical egg/clinical derived HA and NA amino acid sequences. Five strains (4 pH1N1 and 1 H1N2) had between 1-3 amino acid positions in HA that differed between the original clinical material and egg isolate. However, there was no evidence of positive selection for egg adapted variants after a single passage in eggs at any position in HA or NA for either subtype, indicating that the changes observed over time were a result of antigenic drift.

We next examined the observed substitutions in HA and NA in the context of their 3D protein structures. Homology models were constructed using representative sequences of the pH1N1 and huH1N2 strains identified in this study and the locations of the major substitutions mapped. For pH1N1, positions 45 and 83 are surface exposed and located in the globular head of HA1, but not within the five classical antigenic regions (Fig 4B and 4C). Both positions are adjacent to antigenic site Cb, with position 83 close to the conserved N-linked glycosylation site at position 87. Position 345 (HA2 position 18) is located in the fusion peptide, a highly conserved 23-residue sequence responsible for membrane fusion. The NA I10V substitution is located in the N-terminal transmembrane domain (homology models for pH1N1 NA were not produced as the N-terminus is removed by Pronase prior to NA structure determination). For huH1N2, the two major substitution sites in HA are located on the globular head, close to the antigenic regions. Position 172 is buried beneath antigenic site Ca1, whereas position 196 is only partially exposed and located at the top of the globular head, in a pocket close to the junction of antigenic sites Sa and Sb (Fig 4E and 4F). The three major substitution sites in the NA of huH1N2 (313, 385 and 452) are all surface exposed and located on the side of the molecule (Fig 4G). Position 452 is at the bottom of the Mem5 epitope and immediately adjacent to position 253, previously identified in monoclonal antibody escape experiments, whereas positions 313 and 385 are not located in currently mapped antigenic sites [7,39,40].

### Genetic drift of the six internal gene segments

In order to gain a more complete understanding of the evolution of swIAV over time, predicted amino acid sequences from the six internal gene segments of all weaner unit isolates were also aligned and substitutions occurring in the major gene products recorded. Similar to HA and NA, the 6 internal gene segments were relatively static during the first wave of infection (Fig 5). For farm 1, amino acid substitutions in PB2, PB1, PA and NP were detected during this period, however, these were mostly limited to a single sampling time-point and none of the substitutions became fixed (Fig 5A). With the re-emergence of pH1N1 during the second wave of infection, strains containing N350K and K716N in PA displaced previously circulating strains. This strain subsequently acquired M473I in PB2 and R189K in PB1. For huH1N2, amino acid substitutions were observed in all internal gene segments during the first wave of infection, but similar to those observed for pH1N1, most were only detected at a single time-point. An exception was M112I in NS1, which first emerged early in wave 1 but didn’t become fixed until the end of that wave, after which it dominated. Similarly, S82N in M2 emerged at the very end of wave 1 and became fixed thereafter and T257A in PB1 emerged at the start of wave 2, displacing previously circulating variants.

**Fig 5.**
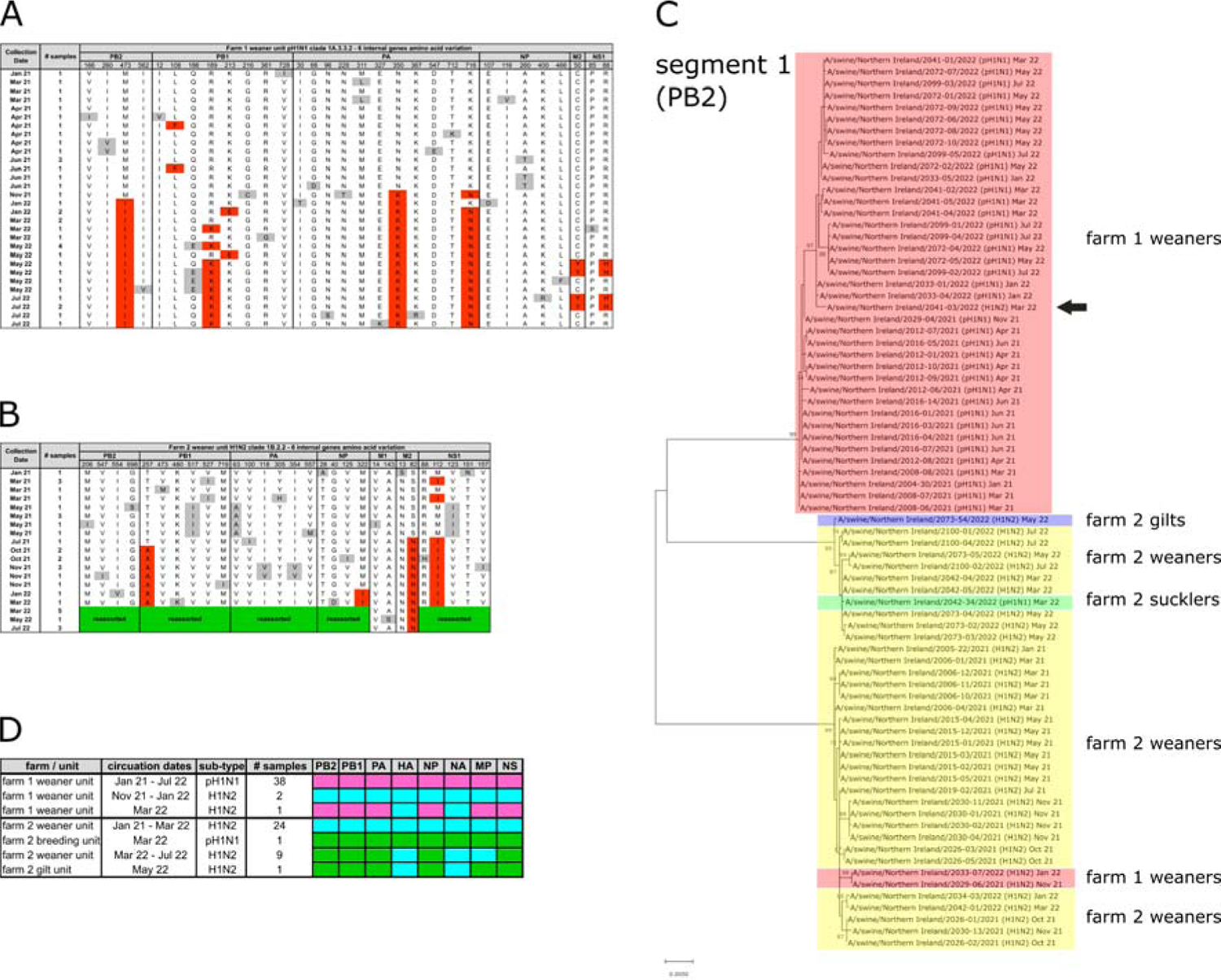
Evolution of the 6-internal gene segments. Amino acid substitutions in the major products of the 6-internal gene segments of (A) farm1 weaner unit pH1N1strains and (B) farm 2 weaner unit huH1N2 strains. Consensus sequences were derived from strains isolated during the first 6 months of the study. Deviations from the consensus are highlighted. Grey highlights indicate substitutions that were detected during a single time-point. Red highlights represent substitutions detected during two or more time-points. Green blocks indicated reassortment of segments. (C) Maximum-Likelihood phylogenetic tree of all pH1N1 and huH1N2 sequences obtained based on complete PB2 coding sequences, with 1,000 bootstrap replications. Trees colour-coded according to the farm/production stage where samples were obtained: farm1 weaners (pink), farm 2 weaners (yellow), farm 2 sucklers (green) and farm 2 gilts (blue). Arrow indicates a reassortant huH1N2, containing internal gene segments from pH1N1. (D) Schematic of the internal gene constellations of all sequenced strains.

### Reassortment of the six internal gene segments

Sequence analysis of huH1N2 isolates obtained from the farm 2 weaning unit indicated that an antigenic shift occurred in March 2022, with a reassortment event involving segments 1, 2, 3, 5 and 8 (Fig 5B). To investigate further, nucleotide sequence alignment of all segments from all available production stages was performed (Fig 5C shows the result of this analysis for segment 1 as an example). This revealed that the five internal gene segments in the reassorted huH1N2 originated from the pH1N1 strain that first emerged in March 2022 on the farm 2 breeding unit and detected in a suckling pig (Fig 5D). On the farm 2 weaning unit, the reassorted huH1N2 displaced the previous strain and was the only strain detected for the remainder of the study, which included the peak of the second infection wave. Interestingly, the partial huH1N2 sequence obtained from a farm 2 gilt in May 2022 contained all six internal gene segments from the pH1N1 strain, indicating the occurrence of a second reassortment event on farm 2.

An independent reassortment event was also detected on farm 1. As described above, pH1N1 was the sole subtype detected on the farm 1 weaning unit from January 2021 until November 2021, when huH1N2 entered the unit and co-circulated until March 2022. Analysis of the six internal gene segments from all farm 1 strains revealed that the internal gene constellation of pH1N1 was unchanged throughout the entire study period. In contrast, a single isolate of huH1N2 obtained in March 2022 was shown to contain the six internal gene segments from the co-circulating pH1N1 (Fig 5C (indicated with arrow) and Fig 5D). huH1N2 was not detected on the farm 1 weaning unit after March 2022.

## Discussion

The study reported here had two major objectives. Firstly, to investigate the infection dynamics of swIAV in intensive pig production systems. Secondly, to characterise the evolution of swIAV in large, infected herds over time. With regard to the first objective, we found that prevalence of swIAV across the different production stages varied considerably. On the breeding units under investigation, prevalence was generally very low. This may be partly explained by the presence of MDA, as levels would be high in piglets under four weeks of age due to sow vaccination. However, studies looking at protection offered by MDA have produced conflicting reports in relation to the extent of protection afforded [41–43]. Respiporc FLU3 does not contain a pH1N1 component and so MDA produced by sows that receive this vaccine would be unlikely to prevent infection of piglets upon exposure. This may partially explain the outbreak of pH1N1 on the farm 2 breeding unit in March 2022. Interestingly, although widespread pH1N1 infection was detected, this was epizootic in nature and limited to a single time-point, clearly indicating that infection dynamics on the breeding units are markedly different from those on weaning units, likely due to the different husbandry practices utilised.

We identify weaned piglets as the major reservoir for continuous circulation of influenza in the intensive farms under investigation. Weaned piglets between 4 and 12 weeks old represent a diverse population in terms of serological status and susceptibility to swIAV. Newly weaned piglets have high levels of MDA due to sow vaccination. However, the efficacy of MDA at this stage is uncertain, with several studies indicating incomplete protection [26,44]. Exposure to swIAV in the presence of MDA has also been associated with impaired immune responses that may facilitate recurrent infections of individual pigs [22]. Despite this, MDA is thought to provide at least partial protection up to ten weeks, therefore progress through this production stage is characterised by waning immunity and increasing susceptibility to infection [45]. The very high prevalence of swIAV on the weaning units suggests that exposure would be frequent and piglets would likely become infected at an early stage post-waning of immunity. Recovered piglets that mount an appropriate humoral response would reduce the pool of susceptible piglets available for infection by homologous strains. However, the high turnover rate – including removal of older exposed pigs to specialised finishing herds and the introduction of newly weaned piglets from the breeding units, means the proportion of pigs susceptible to infection is likely maintained at a level sufficient to sustain continuous infection of the herd.

Measurement of swIAV antibodies in oral fluid samples demonstrated high seroprevalence in finishing pigs in this study. We presume seroconversion occurred following natural infection on the weaning units since swIAV was almost completely absent from both finishing herds for the entire duration of study (except for a single ∼ct 35 detection on farm 2 in May 2022). Introductions into the finishing unit would be expected to occur occasionally but these don’t appear to result in large outbreaks, most likely due to the high degree of pre-existing immunity within the compartment. A previous study of Dutch farrow-to-finish and specialised finishing herds found that swIAV incidences (based on seroconversion per period of time) mostly occurred at the start of the finishing period (12-16 weeks) on farrow-to-finish farms and at the end of the finishing period (16-22 weeks) on specialised finishing herds [18]. This contrasts with our findings and may be explained by lower levels of circulating virus in the preceding production stages of the Dutch farms, resulting in a higher proportion of unexposed/susceptible animals entering the finishing units.

The second objective of this study was to characterise the evolution of swIAV over time in continuously infected intensive herds. Few studies have utilised WGS to study on-farm evolution of swIAV over extended periods. A yearlong study of five farrow-to-wean farms in the United States sequenced 123 complete genomes, identifying 3 subtypes that clustered into 7 distinct viral groups and revealed a complex picture of viral emergence, persistence and subsidence [23]. The same group also sequenced 92 complete genomes during a 15-week cohort study on a wean-to-finish farm, identifying 2 subtypes that clustered into 3 distinct viral groups that co-circulated at different proportions over time [24]. These two studies differ from the findings in the present study, where only a single subtype persisted for the entire duration of study on each farm. Mapping the pattern of amino acid substitutions in the viral surface glycoproteins on the weaner units over time revealed long periods of stasis punctuated by periods of rapid evolution. Such periods of rapid evolution occurred following peaks in swIAV antibody levels in the herd and dramatically reduced viral prevalence. We speculate that periodic reductions in the pool of susceptible animals below maintenance levels, caused by high rates of infection that outpace recruitment of susceptible animals, exerts immune pressure that drives antigenic drift. The location of several of the substitutions noted in HA and NA are suggestive of antigenic drift. However, antigenic characterisation using hemagglutination / neuraminidase inhibition assays will be necessary to demonstrate this conclusively.

In addition to substitutions in the surface glycoproteins, we also detected changes in the other major viral gene products, suggesting that additional selection pressures beyond humoral immunity to HA and NA are involved. Most of these changes emerged in parallel with those in HA and NA and may be epistatic in nature. We also detected a number of reassortant events between established and newly introduced strains. Whilst some reassortant viruses were only detected once, others rapidly displaced strains containing the original internal gene constellation, indicating increased allelic fitness of the new genetic segments. Taken together, these observations suggest a dynamic situation whereby a single strain can persist for extended periods of time without significant antigenic evolution but can rapidly adapt once immune pressure increases due to flux in the size of the susceptible pool. Adaptation is via accumulation of amino acid substitutions in both the surface glycoprotein and the internal gene products, and via exchange of internal gene segments from newly introduced strains. Further analyses are planned to determine the rate of viral evolution and examine positive and negative selection of genetic variants using modelling approaches.

The findings of the present study may contribute to the design of improved infection control strategies. Alternative vaccination protocols that include weaned pigs should be explored to determine the optimum strategy for reducing swIAV prevalence in intensive pig production systems. Furthermore, this study highlights the importance of swIAV monitoring, including genetic characterisation, in order to select the most appropriate vaccines for deployment on farm. In addition, the use of the alternative sampling methodologies described herein should facilitate expanded routine surveillance to monitor herds for the emergence of novel variants, including those with increased zoonotic potential.

## Supporting Information

Supplementary Table S1. Primers utilised for amplicon-based Illumina sequencing.

Supplementary Fig S1. Comparison of viral prevalence with antibody status.

Supplementary Table S2. Comparison of swIAV HA and NA amino acid sequences from clinical material and egg passage 1 (E1) isolates.

## Funding Statement

KL received funding from the Department of Agriculture, Environment and Rural Affairs (DAERA; https://www.daera-ni.gov.uk/) under The Evidence & Innovation programme (Project # 19/3/01). DAERA played no role in the study design, data collection and analysis, decision to publish or preparation of the manuscript.

## Data availability

Sequence data generated in this study have been shared via GISAID, the global data science initiative under the accession numbers EPI_ISL_18221301 to EPI_ISL_18221395.

## Notes

### Competing Interest Statement

The authors have declared no competing interest.

### Summary of Updates

Typo in Fig 2 corrected. Typo in Fig 2 legend corrected. Unlinked reference removed from Discussion text.

