## Supporting Information for "Swine influenza A virus infection dynamics and evolution in intensive pig production systems"

**Supplementary Table S1. Primers utilised for amplicon-based Illumina sequencing.**

| **Influenza A Whole Genome Sequence by Multiplex Primer Assay** | |
| --- | --- |
| HFor Adapter with MBTUni-12 primers | 5' TCG TCG GCA GCG TCA GAT GTG TAT AAG AGA CAG **AGC AAA AGC AGG** |
| HFor Adapter truncated with MBTUni-12 primers | 5' TGT ATA AGA GAC AG**A GCA AAA GCA GG** |
| HRev Adapter with MBTUni-13 primers | 5' GTC TCG TGG GCT CGG AGA TGT GTA TAA GAG ACA G**AG TAG AAA CAA GG** |
| HRev Adapter truncated with MBTUni-13 primers | 5' TGT ATA AGA GAC AG**A GTA GAA ACA AGG** |
| PB2 HFor Adapter | 5’ TCG TCG GCA GCG TCA GAT GTG TAT AAG AGA CAG **AGC RAA AGC AGG TCA AAT** |
| PB2 HFor Adapter truncated | 5' TGT ATA AGA GAC AG**A GCR AAA GCA GGT CAA AT** |
| PB2 HRev adapter | 5' GTC TCG TGG GCT CGG AGA TGT GTA TAA GAG ACA G**AG TAG AAA CAA GGT CGT T** |
| PB2 HRev Adapter truncated | 5' TGT ATA AGA GAC AG**A GTA GAA ACA AGG TCGTT** |
| PB1 HFor Adapter | 5’ TCG TCG GCA GCG TCA GAT GTG TAT AAG AGA CAG **AGC RAA AGC AGG** **CAA ACC** |
| PB1 HFor Adapter truncated | 5' TG TAT AAG AGA CAG **AGC RAA AGC AGG CAA ACC** |
| PB1 HRev Adapter | 5' GTC TCG TGG GCT CGG AGA TGT GTA TAA GAG ACA G**AG TAG AAA CAA GGC ATT T** |
| PB1 HRev Adapter truncated | 5' TGT ATA AGA GAC AG**A GTA GAA ACA AGG CAT TT** |
| PA HFor Adapter | 5’ TCG TCG GCA GCG TCA GAT GTG TAT AAG AGA CAG **AGC RAA AGC AGG** **TAC TGA** |
| PA HFor Adapter truncated | 5' TGT ATA AGA GAC AG**A GCR AAA GCA GGT ACT GA** |
| PA HRev Adapter | 5' GTC TCG TGG GCT CGG AGA TGT GTA TAA GAG ACA G**AG TAG AAA CAA GGT ACT T** |
| PA HRev Adapter truncated | 5' TGT ATA AGA GAC AG**A GTA GAA ACA AGG** **TAC TT** |

Plain font - adapter sequences, Bold font – conserved termini

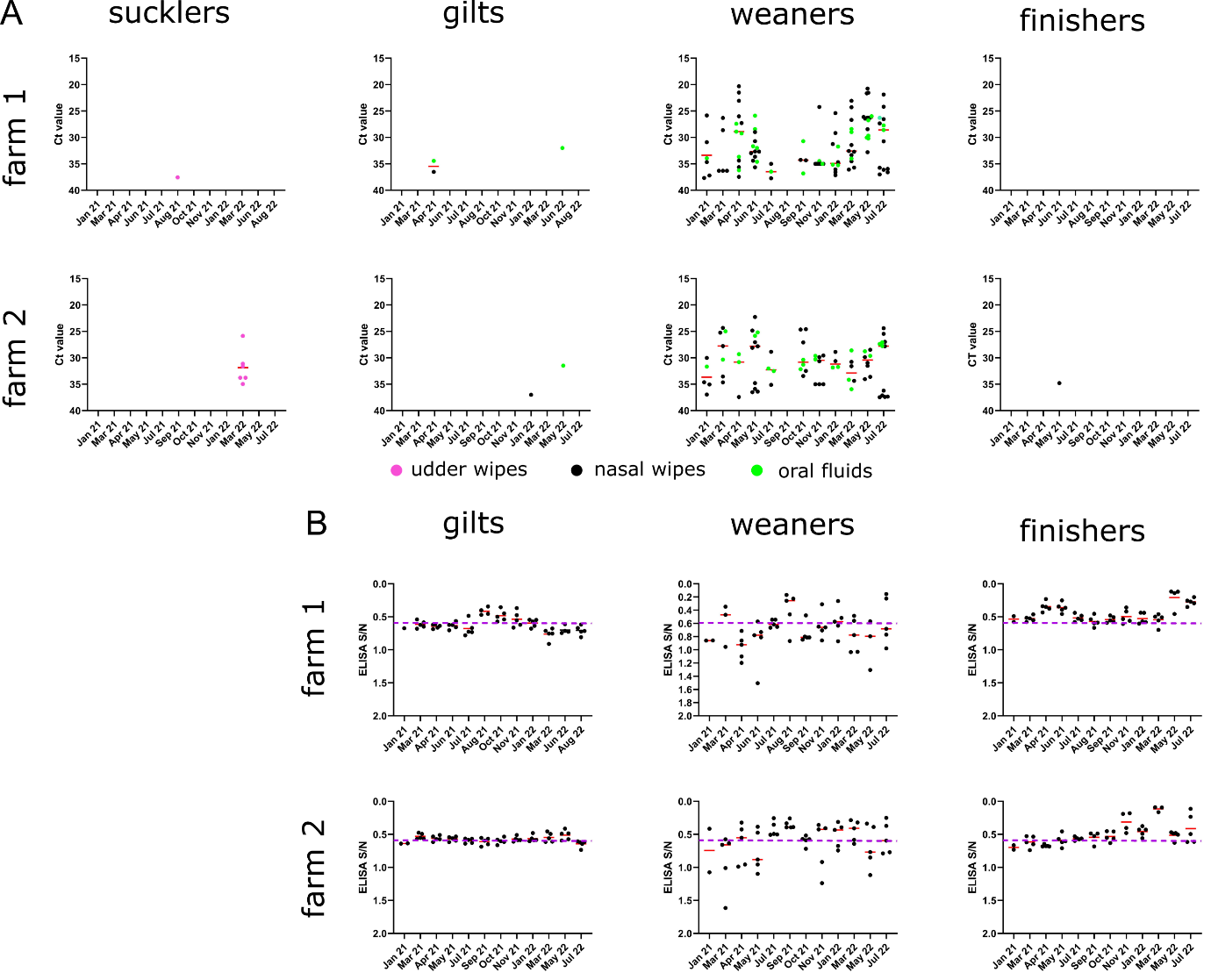

**Supplementary Fig S1. Comparison of viral prevalence with antibody status.** (A) Nasal wipes, udder wipes and oral fluids were tested for the presence of swIAV by RRT-PCR and plotted as individual ct values. Data points coloured according to sample type: udder wipes (pink), nasal wipes (black), oral fluids (green), with mean values indicated by red bars. (B) Oral fluid samples were tested for the presence of influenza NP antibodies by a blocking ELISA and plotted as individual S/N ratios (black circles) with mean values indicated by red bars. S/N ratios ≤0.6 are considered antibody positive. The dashed purple line indicates the 0.6 S/N ratio cut-off value.

**Supplementary Table S2. Comparison of swIAV HA and NA amino acid sequences from clinical material and egg passage 1 (E1) isolates.**

| **Collection Date** | **Farm / Unit** | **Isolate name** | **Sub-type** | **passage** | **HA egg / clinical consensus disagreement** | **NA egg / clinical consensus disagreement** |
| --- | --- | --- | --- | --- | --- | --- |
| Jan 21 | farm 1 weaner unit | A/swine/Northern Ireland/2004-30/2021 | pH1N1 | E1 | E195 | none |
|  |  |  |  | clinical | A195 |  |
| Mar 21 | farm 1 weaner unit | A/swine/Northern Ireland/2008-07/2021 | pH1N1 | E1 | N194 | none |
|  |  |  |  | clinical | D194 |  |
| Apr 21 | farm 1 weaner unit | A/swine/Northern Ireland/2012-06/2021 | pH1N1 | E1 | N35, E127, R509 | none |
|  |  |  |  | clinical | D35, D127, K509 |  |
| Apr 21 | farm 1 weaner unit | A/swine/Northern Ireland/2012-08/2021 | pH1N1 | E1 | none | none |
|  |  |  |  | clinical |  |  |
| Apr 21 | farm 1 weaner unit | A/swine/Northern Ireland/2012-09/2021 | pH1N1 | E1 | none | none |
|  |  |  |  | clinical |  |  |
| Apr 21 | farm 1 weaner unit | A/swine/Northern Ireland/2012-10/2021 | pH1N1 | E1 | none | none |
|  |  |  |  | clinical |  |  |
| Jun 21 | farm 1 weaner unit | A/swine/Northern Ireland/2016-14/2021 | pH1N1 | E1 | none | none |
|  |  |  |  | clinical |  |  |
| Jan 22 | farm 1 weaner unit | A/swine/Northern Ireland/2033-01/2022 | pH1N1 | E1 | N45, N130, I166 | none |
|  |  |  |  | clinical | S45, K130, T166 |  |
| Nov 21 | farm 1 weaner unit | A/swine/Northern Ireland/2029-06/2021 | H1N2 | E1 | none | none |
|  |  |  |  | clinical |  |  |
| Mar 21 | farm 2 weaner unit | A/swine/Northern Ireland/2006-01/2021 | H1N2 | E1 | none | none |
|  |  |  |  | clinical |  |  |
| Mar 21 | farm 2 weaner unit | A/swine/Northern Ireland/2006-04/2021 | H1N2 | E1 | none | none |
|  |  |  |  | clinical |  |  |
| Mar 21 | farm 2 weaner unit | A/swine/Northern Ireland/2006-11/2021 | H1N2 | E1 | V30, V309 | none |
|  |  |  |  | clinical | I30, A309 |  |
| May 21 | farm 2 weaner unit | A/swine/Northern Ireland/2015-01/2021 | H1N2 | E1 | none | none |
|  |  |  |  | clinical |  |  |
| May 21 | farm 2 weaner unit | A/swine/Northern Ireland/2015-02/2021 | H1N2 | E1 | none | none |
|  |  |  |  | clinical |  |  |
| May 21 | farm 2 weaner unit | A/swine/Northern Ireland/2015-03/2021 | H1N2 | E1 | none | none |
|  |  |  |  | clinical |  |  |
| May 21 | farm 2 weaner unit | A/swine/Northern Ireland/2015-12//2021 | H1N2 | E1 | none | none |
|  |  |  |  | clinical |  |  |
| Oct 21 | farm 2 weaner unit | A/swine/Northern Ireland/2026-02/2021 | H1N2 | E1 | none | none |
|  |  |  |  | clinical |  |  |
| Oct 21 | farm 2 weaner unit | A/swine/Northern Ireland/2026-05/2021 | H1N2 | E1 | none | none |
|  |  |  |  | clinical |  |  |
